# Perception of invisible masked objects in early infancy

**DOI:** 10.1101/2021.02.15.431195

**Authors:** Yusuke Nakashima, So Kanazawa, Masami K. Yamaguchi

## Abstract

Recurrent loops in the visual cortex play a critical role in visual perception, which is likely not mediated by purely feedforward pathways. However, the development of recurrent loops is poorly understood. The role of recurrent processing has been studied using visual backward masking, a perceptual phenomenon in which a visual stimulus is rendered invisible by a following mask, possibly because of the disruption of recurrent processing. Anatomical studies have reported that recurrent pathways are immature in early infancy. This raises the possibility that younger infants process visual information mainly in a feedforward manner, and thus, they might be able to perceive visual stimuli that adults cannot see because of backward masking. Here, we show that infants under 7 months of age are immune to visual backward masking and that masked stimuli remain visible to younger infants while older infants cannot perceive them. These results suggest that recurrent processing is immature in infants under 7 months and that they are able to perceive objects even without recurrent processing. Our findings indicate that the algorithm for visual perception drastically changes in the second half of the first year of life.

## Introduction

The standard view of cortical visual processing is that visual information is hierarchically processed along feedforward pathways, with more complex representations created serially by relaying information from lower to higher visual areas (1). However, recurrent processing, which is mediated by corticocortical feedback and intra-areal horizontal pathways (2, 3), also contributes to fundamental visual functions (4*-*6). Although the role of recurrent processing is not yet well understood, many studies have proposed that visual perception is not mediated by a purely feedforward system but rather by a system incorporating recurrent loops (7*-*12).

Anatomical studies of the infant brain have reported that feedback and horizontal connections develop later than feedforward connections (13-15). Anatomical data from postmortem brains of human infants have shown that the adult-like laminar termination pattern of forward connections between V1 and V2 emerges by 4 months of age (14), but the feedback (14) and long-range horizontal (15) connections are immature at that age. These findings imply that until at least around the second half of the first year of life, recurrent processing is immature, and visual information may be processed mainly in a feedforward manner. However, this possibility has so far not been examined. Although neuroimaging studies of human infants have revealed functional and structural brain development (16, 17), it is difficult to determine whether the observed activities or structures are derived from feedforward or recurrent pathways using imaging techniques in human infant studies. Thus, in the present study, we examined recurrent processing in early infancy using visual backward masking.

Visual backward masking is a perceptual phenomenon in which a stimulus is rendered invisible by a mask presented after the target stimulus. We used object substitution masking (OSM) (18), a type of backward masking, which is thought to arise from a disruption of recurrent processing (18*-*23). In OSM, target perception is impaired when a target is briefly presented with a sparse mask that surrounds the target and the mask remains on-screen after the target disappears, while target perception remains intact when the target and the mask disappear simultaneously. OSM has been proposed to occur because the temporally trailing mask disrupts the recurrent signals related to the target (18). Although some studies have questioned the recurrent explanation (24, 25), evidence from psychophysical studies suggests that OSM can be plausibly explained by the recurrent theory rather than the feedforward theory (18, 20, 22, 23). Indeed, a neuroimaging study has shown that recurrent activities in the early visual cortex are modulated when target perception is impaired by OSM (19).

Although an infant study has suggested that OSM occurs in 6-month-old infants (26), its mechanism and development remain poorly understood. If visual processing is performed without recurrent processing in early infancy, as suggested by the anatomical studies (14, 15), OSM may not occur in younger infants. In other words, younger infants may be able to perceive a masked stimulus that older infants cannot.

## Results

### OSM is present in older infants

We first examined the development of OSM in 3-to 8-month-old infants, using a preferential looking paradigm with face stimuli (Fig. 1A). It is known that infants show spontaneous preferences for faces over non-face objects (27). Thus, by measuring preferences for a face, we can test whether or not infants can perceive the face followed by a mask. In an adult study, OSM has been shown to occur when faces are used as a target (28).

**Fig 1.**
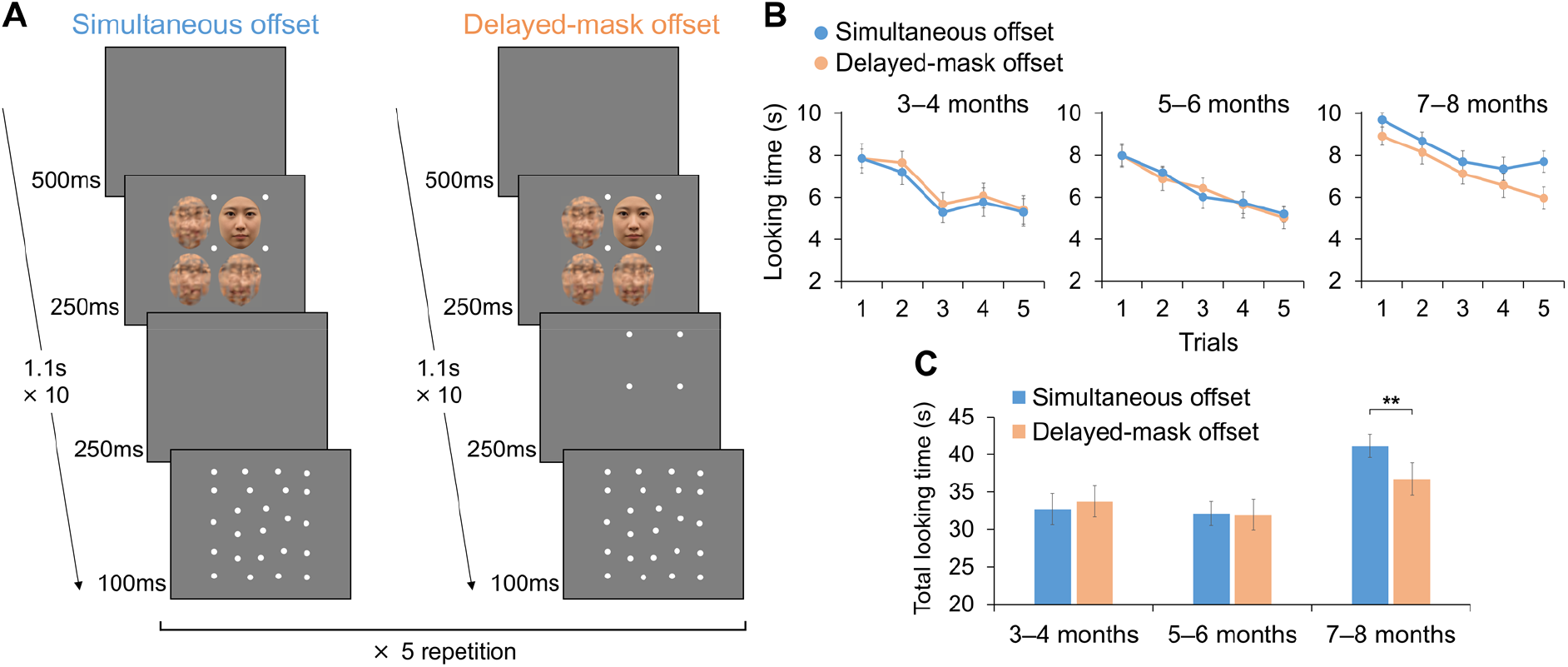
OSM occurs in 7- to 8-month-old infants. (**A**) The two conditions in the first experiment. If OSM occurs, faces can be perceived in the simultaneous offset condition but not in the delayed-mask offset condition. Thus, looking times should be longer in the simultaneous offset condition. (**B**) Looking times for simultaneous offset (blue) and delayed-mask offset (orange) conditions on each trial. (**C**) Total looking times on the five trials for each condition. See Fig. S1A for individual data. All data are mean ± s.e.m. \*\**P* < 0.01.

A face was presented with three distractors (scrambled faces) for 250 ms, during which the face was surrounded by a four-dot mask typically used in OSM (18). In the delayed-mask offset condition (masked condition), the mask remained on the screen for 250 ms after the face and distractors disappeared. In the simultaneous-offset condition (unmasked condition), the face and mask disappeared simultaneously. The stimulus sequence was repeated for 11 s on each trial, and the infant’s looking time at the stimulus array during the trial was measured. Trials in the two conditions were presented alternately and were repeated five times. If OSM occurs, infants should not perceive faces in the delayed-mask offset condition, and looking times at the stimulus array should be longer in the simultaneous offset condition, in which faces can be perceived.

Infants were separated into three age groups: 3–4 months, 5–6 months, and 7–8 months of age. Looking time was different between the two conditions only in 7-to 8-month-old infants (Fig. 1B, C). Total looking time on the five trials was compared between the two conditions (Fig. 1C). Infants at 7–8 months looked longer in the simultaneous-offset condition (paired *t*-test, *t*_19_ = 3.36, p < 0.01, d = 0.52), while infants at 3–4 months and at 5–6 months showed no differences between conditions (3–4 months: *t*_19_ = −1.12, p = 0.56, d = −0.11; 5–6 months: *t*_19_ = 0.19, p = 0.86, d = 0.02). These results indicate that only 7-to 8-month-old infants could not perceive faces in the delayed-mask offset condition, suggesting that OSM begins to occur in 7-to 8-month-old infants.

### OSM is ineffective for younger infants

Although there was no difference between the masked and unmasked conditions in 3-to 6-month-old infants in the first experiment, we cannot conclude that masking does not occur in younger infants based solely on non-significant results. To further test the absence of OSM in younger infants, we next compared delayed-mask offset and no-face conditions (Fig. 2A). The delayed-mask offset condition was same as that in the first experiment, and the no-face condition was the same as the simultaneous-offset condition, except that a face was not presented at the target position. If OSM does *not* occur, infants should perceive faces in the delayed-mask offset condition, and looking time should be longer in the delayed-mask offset condition than in the no-face condition.

**Fig 2.**
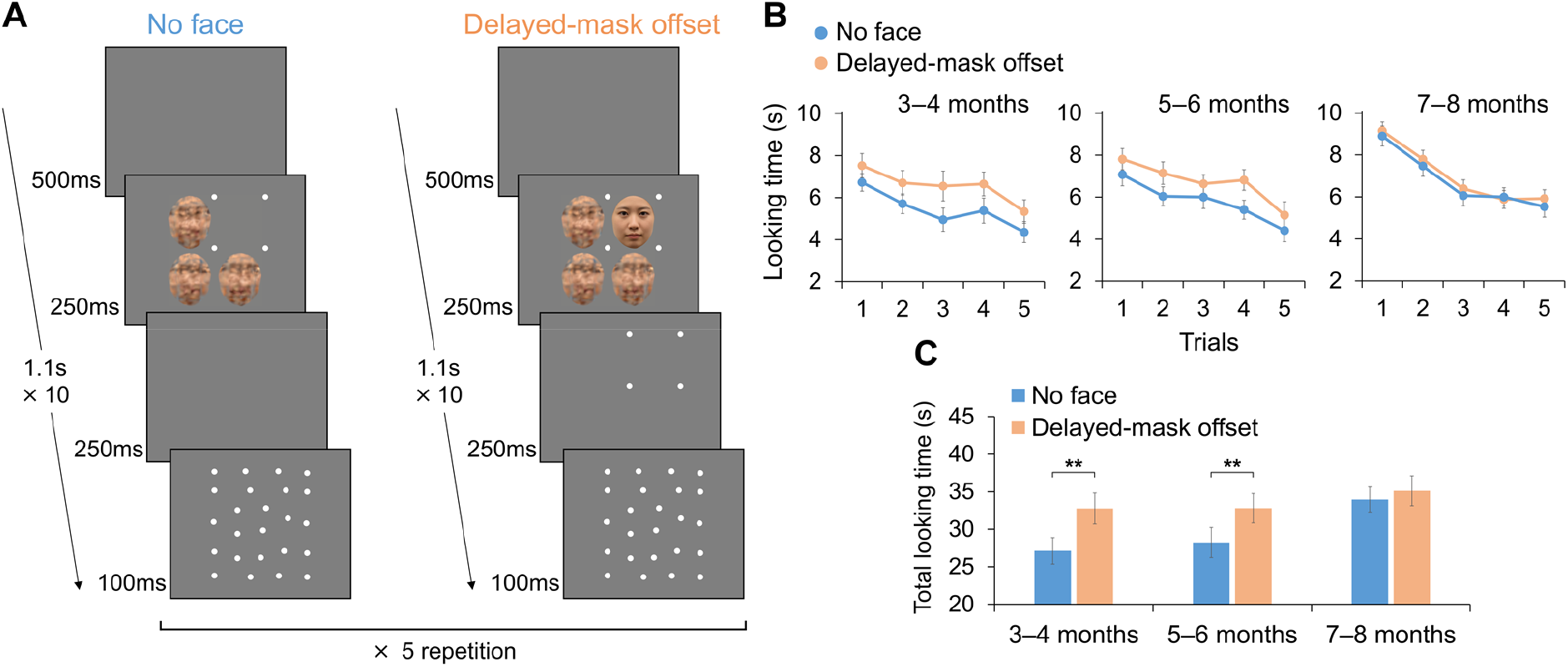
Infants at 3–6 months are immune to OSM and can perceive faces in the masked condition. (**A**) The no-face and delayed-mask offset conditions in the second experiment. If OSM does *not* occur, looking times should be longer in the delayed-mask offset condition. (**B**) Looking times for the no-face (blue) and delayed-mask offset (orange) conditions on each trial. (**C**) Total looking times on the five trials for each condition. See Fig. S1B for individual data. All data are mean ± s.e.m. \*\**P* < 0.01.

In contrast to the first experiment, a difference in looking time between the two conditions was observed for infants at 3–4 months and 5–6 months but not at 7–8 months (Fig. 2B, C). Total looking time on the five trials was longer in the delayed-mask offset condition for infants at 3–4 and 5–6 months (3–4 months: *t*_19_ = 6.15, p < 0.01, d = 0.66; 5–6 months: *t*_19_ = 5.26, p < 0.01, d = 0.52), while there was no difference for 7–8-month-old infants (*t*_19_ = 1.18, p = 0.25, d = 0.14; Fig. 2C). These results indicate that the persisting four-dot mask was not effective for 3-to 6-month-old infants and that they could perceive faces that the older infants could not. Taken together, these results suggest that OSM occurs in infants over 6 months of age as it does in adults, whereas younger infants are immune to OSM.

### OSM is ineffective for younger infants even with a contour mask

Infants under 7 months of age were able to perceive a face in the masked condition. However, the four-dot mask may have been insufficiently strong to cause OSM in younger infants. That is, 3-to 6-month-old infants may have perceived the four-dot mask as four separate dots, rather than as an object (a blank square produced by four dots). To test this possibility, we examined whether the results for younger infants could be replicated with a stronger mask, an elliptical contour, instead of four dots. The delayed-mask offset and no-face conditions were compared (similar to the second experiment) in 5-to 6-month-old infants (Fig. 3A). The delayed-mask offset condition was the same as that in the previous experiments except that the mask was an elliptical ring, and the no-face condition was same as the delayed-mask offset condition except that the face was replaced by a distractor.

**Fig 3.**
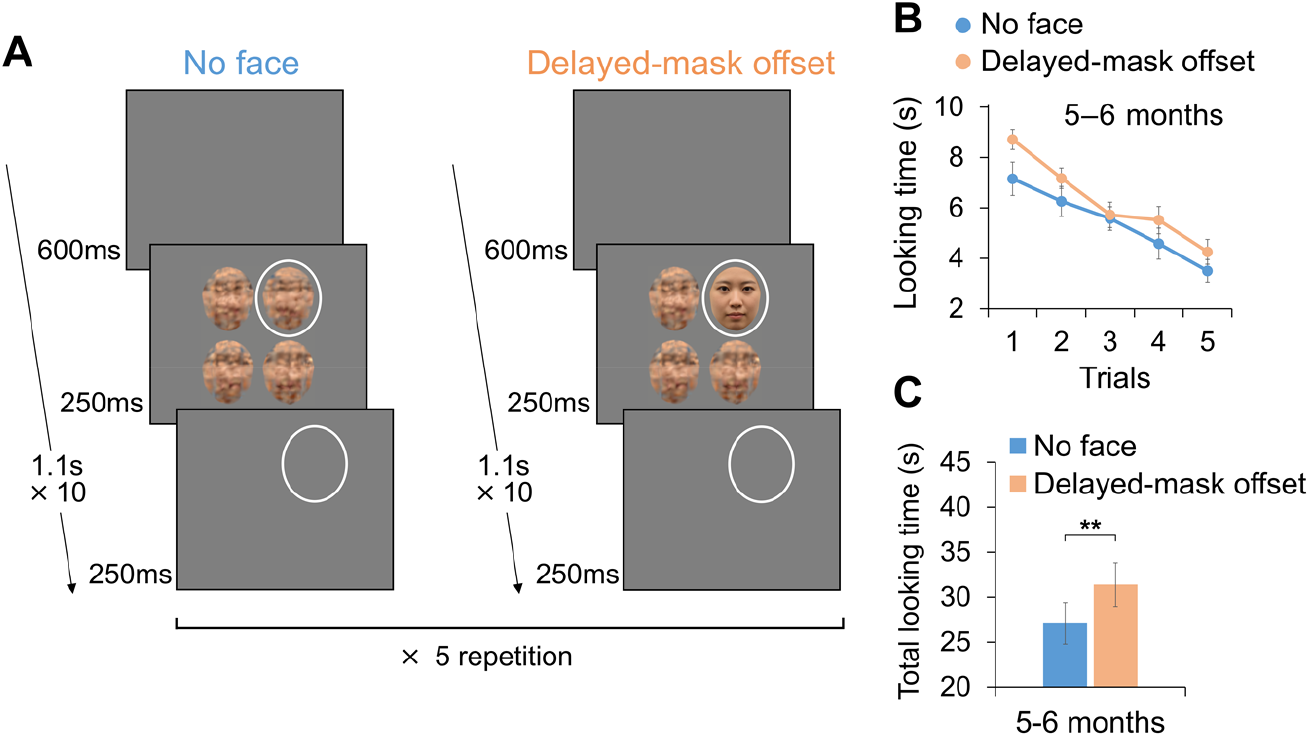
Infants at 5–6 months can perceive faces even with the contour mask. (**A**) In the third experiment, a stronger mask, an elliptical ring, was used in the same conditions as in the second experiment. (**B**) Looking times for the no-face (blue) and delayed-mask offset (orange) conditions on each trial. (**C**) Total looking times on the five trials for each condition. See Fig. S1C for individual data. All data are mean ± s.e.m. \*\**P* < 0.01.

The results replicated those of the second experiment (Fig. 3B, C). Infants at 5–6 months looked longer in the delayed-mask offset condition (*t*_19_ = 3.47, p < 0.01, d = 0.48). Thus, infants at 5–6 months could perceive faces although the mask was stronger, supporting the conclusion that OSM is ineffective for infants under 7 months.

### Validation of the preferential looking paradigm

In these three experiments, it was possible that faces were presented so briefly that infants could not perceive them clearly even without masking. To confirm the validity of our paradigm, we examined whether infants preferred faces without masking. We compared looking times between face and no-face conditions without masks (Fig. S2A, D). The two types of comparisons were conducted in separate experiments, that is, face versus no-face with a blank in the target position (blank experiment) and face versus no-face with a distractor in the target position (distractor experiment).

In the blank experiment, infants in all age groups had longer looking times in the face condition (3–4 months: *t*15 = 3.49, p < 0.01, d = 0.58; 5–6 months: *t*15 = 3.96, p < 0.01, d = 0.57; 7–8 months: *t*15 = 7.49, p < 0.01, d = 0.74; Fig. S2B, C). In the distractor experiment as well, infants in all age groups preferred the face condition (3–4 months: *t*15 = 3.54, p < 0.01, d = 0.52; 5–6 months: *t*15 = 6.51, p < 0.01, d = 0.65; 7–8 months: *t*15 = 4.94, p < 0.01, d = 0.96; Fig. S2E, F). Thus, we confirmed that infants could clearly perceive faces and discriminate them from distractors.

## Discussion

The present study revealed that OSM occurs in 7-to 8-month-old infants but not in 3-to 6-month-old infants. Importantly, the target masked by OSM, which disrupts perception in adults, remained visible to infants at 6 months and younger. This provides evidence of distinct mechanisms of visual perception in younger infants and adults. Given the anatomical studies showing immature recurrent pathways in early infancy (14, 15), the present results could be interpreted as suggesting that recurrent processing is immature in infants at 6 months and younger, and that they can perceive visual stimuli without mature recurrent processing.

Our results demonstrate that a stimulus that can be perceived in early infancy becomes imperceptible during development. This counterintuitive result suggests that the algorithm for visual perception drastically changes in the second half of the first year. Visual processing may be performed through a feedforward system without (or with immature) recurrent processing until around 6 months, while it is performed through the recurrent loops in older infants and adults. Several positive functions would be acquired by incorporating recurrent processing, but it also may lead to backward masking since recurrent processing becomes critical to perceive objects.

What positive functions are acquired by maturing recurrent loops? Recurrent processing is related to the ability to integrate local information into a global context, such as figure-ground segregation (4, 6), illusory contour (29), and surround suppression (30). Thus, these functions are expected to be lacking in younger infants that do not have mature recurrent processing. Indeed, orientation-based texture segregation develops after 5 months (31), and illusory contours can be robustly perceived after 7 months (32, 33). Surround suppression is also immature until 6 months (34). These complex functions develop later than processing of simple features such as orientation and motion selectivity, which develops by 3 months (35). These facts, together with our results, imply that basic processing such as simple feature selectivity is mediated by feedforward pathways, while more complex functions that require global integration are acquired through the development of the feedback and horizontal connectivity.

Some studies have proposed that OSM can be explained by the disruption of feedforward processing without assuming recurrent feedback (24, 25). However, our results are somewhat difficult to reconcile with the feedforward theory of OSM. A feedforward account of the absence of OSM in younger infants may posit that the forward processing of a target is so fast that a masker does not catch up with it. It is unlikely, however, that forward processing is faster in younger infants than in older infants. Myelination in the occipital lobe is immature in infants (36), and the latencies of event-related potentials associated with visual perception are longer in infants than in adults (37, 38). Thus, in line with other studies suggesting that OSM is involved in recurrent processing (18-23), the recurrent account is a more plausible explanation for our results.

Another important function related to recurrent loops is predictive processing. Theoretical studies assume that the visual system constantly integrates top-down predictions with sensory inputs to resolve perceptual ambiguity (7, 11, 12). Recurrent processing has been shown to be important for robustly perceiving degraded stimuli such as occluded or unclear images (39, 40). In early infancy, such predictive mechanisms may be immature, and perception could be vulnerable to degradation and ambiguity. Perhaps, in return for susceptibility to visual masking, robustness of visual perception is acquired. On the other hand, it has been shown that visual cortical activity is modulated by top-down predictions in infants at 6 and 12 months (41, 42). These studies used association learning tasks with visual and auditory stimuli; thus, they were related to cross-modal processing. Moreover, association learning tasks may be related to the subcortical pathways (43). We infer that the mechanisms related to those tasks are different from those in the present study and that their developmental time course may also be different.

## Materials and Methods

### Participants

Sixty infants participated in the first experiment. The infants were divided into three age groups: 3-to 4-month-olds (8 female and 12 male, age range 90–132 days, mean age 115.5 days), 5-to 6-month-olds (9 female and 11 male, age range 148–191 days, mean age 170.8 days), and 7-to 8-month-olds (12 female and 8 male, age range 207–254 days, mean age 233.2 days). An additional 17 infants were excluded from the analysis because of fussiness (n = 10) or insufficient looking time (n = 7; the total looking time on the ten trials was less than 30% of the entire duration of the trials).

Sixty infants participated in the second experiment and were divided into three age groups: 3-to 4-month-olds (11 female and 9 male, age range 82–133 days, mean age 109.0 days), 5-to 6-month-olds (11 female and 9 male, age range 150–194 days, mean age 170.5 days), and 7-to 8-month-olds (10 female and 10 male, age range 204–254 days, mean age 227.9 days). An additional 16 infants were excluded because of fussiness (n = 8) or insufficient looking time (n = 8).

Twenty infants at 5–6 months participated in the third experiment (7 female and 13 male, age range 136–190 days, mean age 162.1 days). An additional infant was excluded because of insufficient looking time.

In the fourth experiments, 96 infants participated. In the blank experiment, 48 infants were divided into three age groups: 3-to 4-month-olds (9 female and 7 male, age range 86–134 days, mean age 113.3 days), 5-to 6-month-olds (9 female and 7 male, age range 142–194 days, mean age 171.8 days), and 7-to 8-month-olds (11 female and 5 male, age range 204–254 days, mean age 230.8 days). An additional nine infants were excluded because of fussiness (n = 6) or insufficient looking time (n = 3). In the distractor experiment, 48 infants were divided into three age groups: 3-to 4-month-olds (7 female and 9 male, age range 85–134 days, mean age 114.5 days), 5-to 6-month-olds (9 female and 7 male, age range 152–194 days, mean age 172.8 days), and 7-to 8-month-olds (11 female and 5 male, age range 200–253 days, mean age 230.5 days). An additional six infants were excluded because of fussiness (n = 5) or insufficient looking time (n = 1).

The infants were recruited through newspaper advertisements. All infants were at full term at birth and healthy at the time of the experiment. This study was conducted according to the Declaration of Helsinki and was approved by the Ethical Committee of Chuo University. Written informed consent was obtained from parents.

### Apparatus and visual stimuli

Visual stimuli were presented on an LCD monitor (Cambridge Research Systems, Display++, 32 inch, 1920 × 1080 resolution, 120 Hz) using PsychoPy 1.90.3. An infant sat on a parent’s lap in front of the display at a viewing distance of 60 cm in a dark room. A camera was set below the display to monitor and record the infant’s looking behavior. An experimenter observed the infant’s behavior through a monitor connected to the camera.

Face stimuli were created by morphing photo images of female frontal view faces. Five face stimuli were created; 4–8 different persons were morphed for each face stimulus. All face stimuli were cropped into an identical shape (5.3° in width, 6.6° in height) to remove the outer features. Distractors were created by local-phase scrambling of a face: the face image was separated into nine square areas and phase-scrambling was done in each area. The scrambled image was then cropped and a Gaussian blur was applied. Six scrambled images were created, and three were randomly chosen as distractors for each trial. The size and contour of distractors were the same as the face stimuli. A face and three distractors were presented in the cells of a 2 × 2 matrix on a uniform gray background (50.2 cd/m^2^). The distance between the centers of the monitor and the stimuli was 3.2° on the horizontal axis and 3.4° on the vertical axis.

The four-dot mask consisted of four white dots (0.7° in diameter, 98.8 cd/m^2^) located at the corners of the face stimulus. The distance between each dot and the center of the face stimulus was 3.1° on the horizontal and vertical axes. In the third experiment, the ring mask (6.6° in width, 7.8° in height, 0.3° in line width, 98.8 cd/m^2^) was presented instead of the four-dot mask. The spacing between the ring mask and the contour of the faces was at least 0.4°. The dot array presented after the four-dot mask consisted of a few dozen dots (each identical to those in the four-dot mask) and subtended 12.8° (width) by 15.5° (height).

### Experimental procedure

We used the preferential looking paradigm to test whether infants could perceive faces. Trials of two conditions were presented sequentially, and we measured infants’ looking behavior to examine which condition infants looked at longer.

In the first experiment, the delayed-mask offset and simultaneous-offset conditions were compared in 3-to 8-month-old infants (Fig. 1A). A face surrounded by the four-dot mask and three distractors were presented for 250 ms. In the delayed-mask offset condition, the mask remained on the screen for 250 ms after the face and distractors disappeared. In the simultaneous-offset condition, the face and mask disappeared simultaneously and were followed by a 250-ms blank. The dot array was then presented for 100 ms in both conditions to prevent the persisting four-dot mask from attracting the infants’ attention. The stimulus sequence was repeated ten times at 500-ms intervals (i.e., 11 s) in a trial. On each repetition, the face and distractors were located at random positions in the 2 × 2 matrix, but a face did not appear in the same position sequentially so that infants did not see the face in central vision. Five trials were presented in each of the two conditions. The same face was presented ten times within each of the five trials, and five different faces were used on the five trials. The ordering of the five face stimuli (i.e., ordering of the trials) within each condition was randomized for each infant. Trials in the two conditions were presented alternately, with the order of the two conditions counterbalanced across infants. Each trial started when the infant looked at a cartoon presented at the center of the monitor at the same time as a short beeping sound. The infant’s looking time at the stimulus array during the trial was measured. If OSM occurred, the looking time would be longer in the simultaneous-offset condition.

In the second experiment, the delayed-mask offset and no-face conditions were compared in 3-to 8-month-old infants (Fig. 2A). The delayed-mask offset condition was the same as that in the first experiment. The no-face condition was the same as the simultaneous-offset condition except that the face was replaced by a blank. If OSM did *not* occur, the looking time would be longer in the delayed-mask offset condition.

In the third experiment, the delayed-mask offset and no-face conditions were compared in 5-to 6-month-old infants (Fig. 3A). The same two conditions as in the second experiment were compared, but an elliptical ring was used as the mask instead of the four dots. In the no-face condition, unlike that in the second experiment in which a blank was surrounded by the mask, a distractor was surrounded by the mask. Also, unlike the previous no-face condition, the mask remained on-screen after the target disappeared to exclude the possibility that infants would prefer the persisting mask. This enabled a fairer comparison of the delayed-mask offset and no-face conditions. The dot array was not presented in this experiment because the mask was presented in both conditions.

In the fourth experiments, to confirm validity of our paradigm, the face and no-face conditions were compared without the mask in 3- to 8-month-old infants (Fig. S2, A and B). In the face condition, the face and the four dots coterminated (as in the simultaneous-offset condition of the first experiment). The no-face conditions were same as the face condition except that the face was replaced by either a blank or a distractor. The two experiments were conducted separately: face versus no-face (blank) and face versus no-face (distractor).

### Data coding and analysis

An observer, blind to the stimuli presented on the trials, measured each infant’s looking times from recoded videos. The looking time for each trial was measured by pressing a key while the infant looked at the stimulus array.

Sample size was estimated based on data of previous infant studies that measured looking time (44) and was calculated to achieve a power of 0.8 with an effect size of 0.65 in the first, second, and third experiments (masking paradigm). In the fourth experiments (non-masking paradigm), sample size was determined to achieve a power of 0.8 with an effect size of 0.75.

Paired *t* tests were conducted to compare the total looking times on five trials between the two conditions for each age group. All *t* tests were two-sided. The *p* values were adjusted using the Bonferroni-Holm method for the three comparisons of each age group in all experiments except the third.

## Acknowledgments

We thank S. Tsurumi, J. Yang, Y. Ujiie, K. Sato, and Y. Tsuji for their assistance with data collection and discussions. This study was supported by JSPS KAKENHI grant number 19K14479 (Y.N.) and MEXT KAKENHI grant number 17H06343 (M.K.Y.)

## Author contributions

Y.N., S.K., and M.K.Y. designed the study. Y.N. performed experiments and analyzed data. Y.N., S.K., and M.K.Y. interpreted the data. Y.N. wrote the original manuscript. Y.N., S.K., and M.K.Y. reviewed and revised the manuscript.

## Data availability

The data used to generate the findings of this study are available in “figshare” with the identifier “doi: 10.6084/m9.figshare.13220873”.

## Supplementary Materials

**Fig S1.**
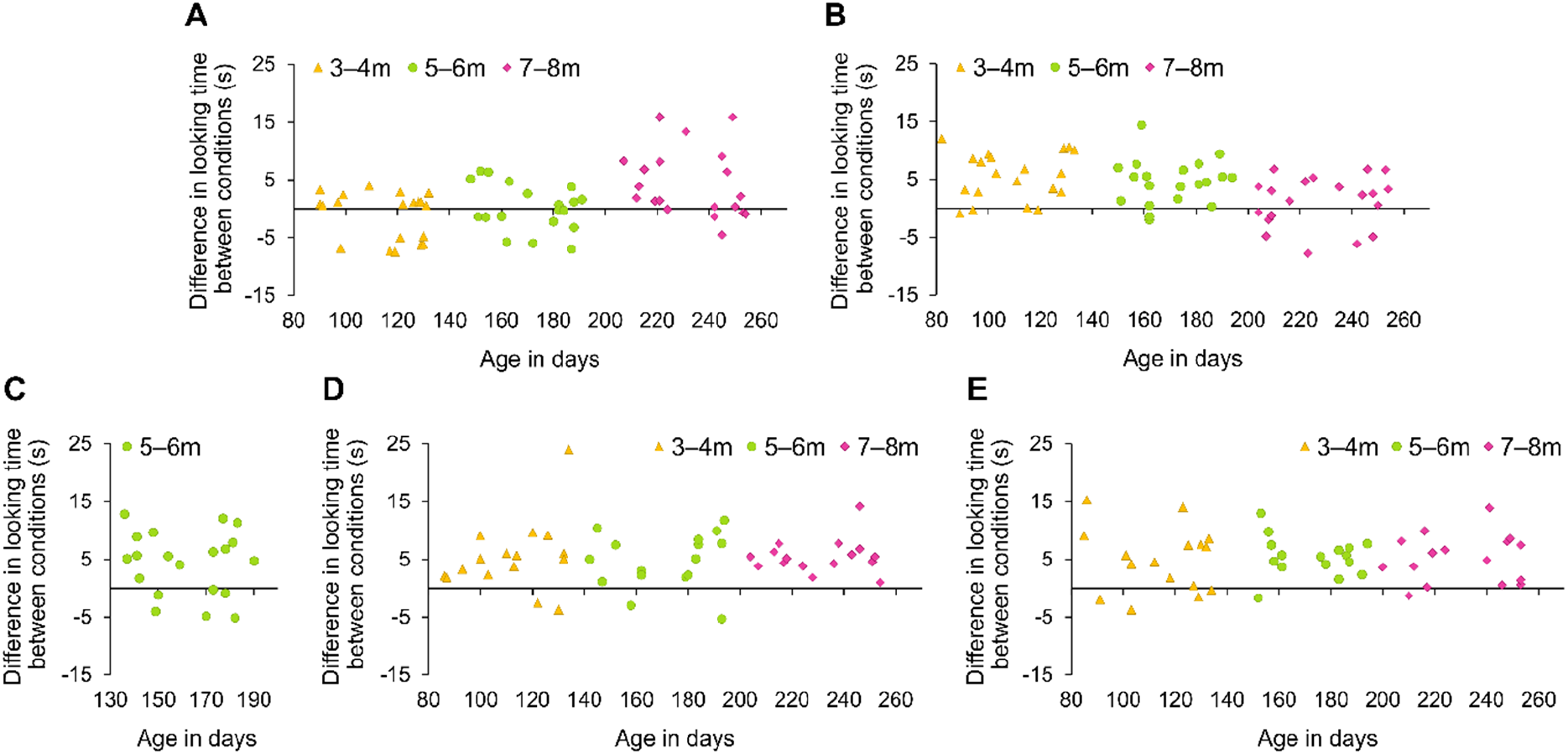
Individual data for each experiment. Differences in total looking time between the two conditions as a function of the infants’ age in days. (**A**) Data from the first experiment, calculated by subtracting looking time in the delayed-mask offset from that in the simultaneous offset condition. (**B**) Data from the second experiment, calculated by subtracting looking time in the no-face condition from that in the delayed-mask offset condition. (**C**) Data from the third experiment, calculated by subtracting looking time in the no-face condition from that in the delayed-mask offset condition. (**D, E**) Data from the fourth experiments, calculated by subtracting looking time in the no-face condition from that in the face condition, for the blank (d) and distractor (e) experiments. Yellow triangles, 3–4 months; green circles, 5–6 months; purple diamonds, 7–8 months.

**Fig S2.**
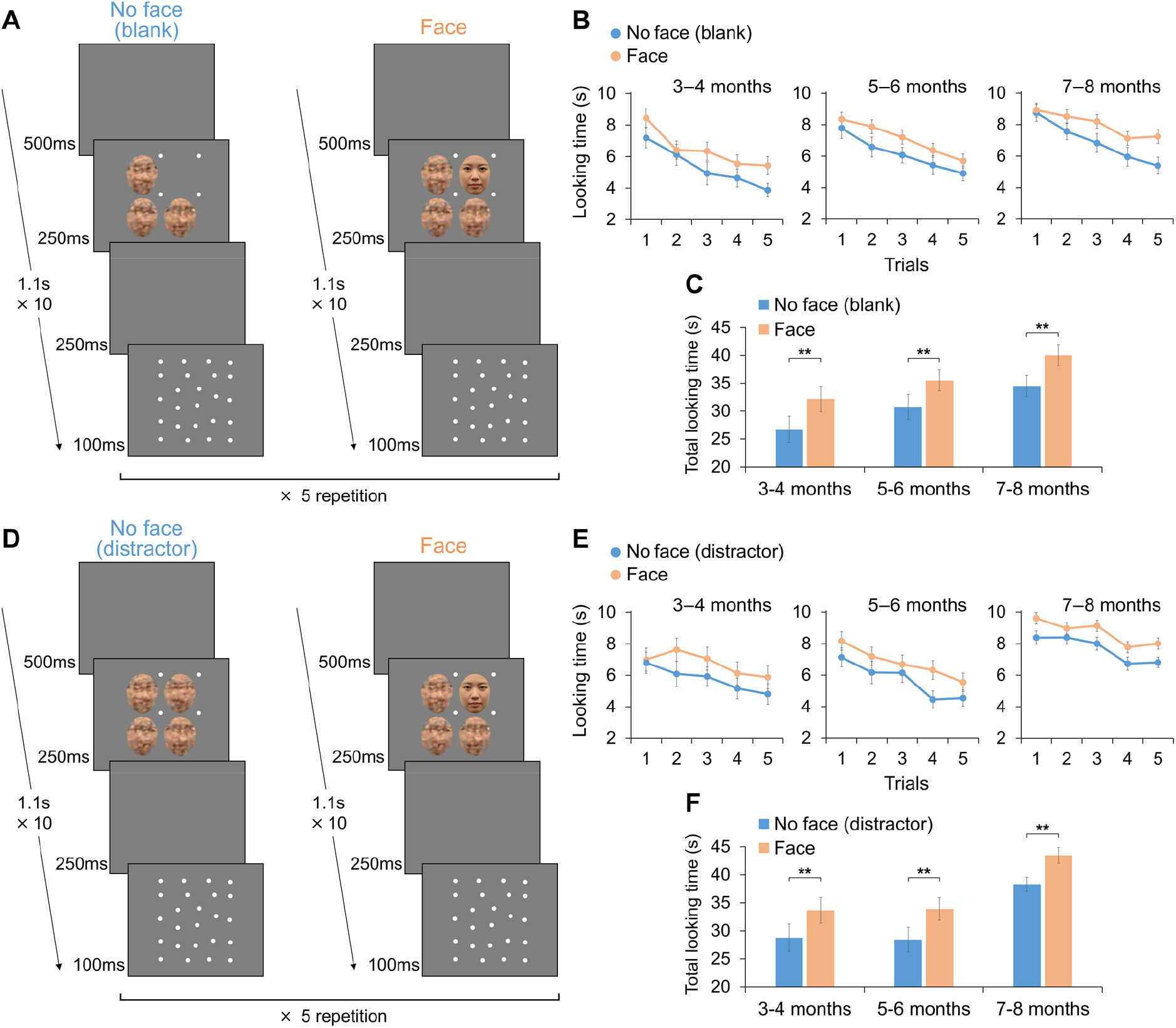
Procedures and results in the fourth experiments. (**A**) No-face (blank) and face conditions without masking. (**B, E**) Looking times for the no face (blue) and face (orange) conditions on each trial. (**C, F**) Total looking times on the five trials for each condition. See Fig. S1D, E for individual data. (**D**) No-face (distractor) versus face conditions. All data are mean ± s.e.m. \*\**P* < 0.01.

